# How a tarantula can help treat infections: *Avicularia juruensis*’s theraphotoxins that also act as antimicrobials

**DOI:** 10.1101/2022.10.09.511474

**Authors:** Soraia Maria do Nascimento, Andrea Díaz Roa, Ronaldo Zucatelli Mendonça, Pedro Ismael da Silva Junior

**Affiliations:** Laboratory of Applied Toxinology - Center of Toxins, Immune-Response and Cell Signaling (CeTICS/CEPID), Butantan Institute, São Paulo, Brazil; Postgraduate Program Interunits in Biotechnology, USP/IBu/IPT, São Paulo, Brazil; Laboratory of Parasitology, Butantan Institute, São Paulo, Brazil

**Keywords:** venom, spider, antimicrobial, Theraphosidae, *Avicularia juruensis*

## Abstract

Considering that there are still many species little-studied, this work aimed to analyze the venom of the spider *Avicularia juruensis* searching for antimicrobial peptides. Using reverse-phase high-performance liquid chromatography, microbial growth inhibition assay, transcriptomics, and proteomics approaches we identified three antimicrobial peptides: Avilin, Juruin_2, and Juruenine. All of them showed similarities with neurotoxins that act on ion channels and, probably, they have the ICK motif. The study of animal venoms is of great importance to carry out the characterization of unknown components and that may have a biotechnological application, in special venoms from spiders that are from less studied families.

Spiders are the most successful group of venomous animals, comprising more than 50,350 species distributed in all terrestrial habitats. One strategy that facility their broad distribution is the production of elaborate venoms, which are composed of inorganic salts, organic molecules with low molecular mass, free amino acids, small polypeptides, linear peptides, nucleotides, disulfide-rich peptides, enzymes, and high molecular mass proteins. Considering that there are still many species little-studied, this work aimed to analyze the venom of the mygalomorph spider *Avicularia juruensis* searching for new antimicrobial peptides. Using reverse-phase high-performance liquid chromatography, microbial growth inhibition assay, transcriptomics, and proteomics approaches we identified three antimicrobial peptides that were named Avilin, Juruin_2, and Juruenine. All of them showed similarities with neurotoxins that act on ion channels and, probably, they have the ICK motif in their structure. The ICK fold seems to be conserved in several venomous animal lineages and presents elevated functional diversity, as well as gives stability to the toxins. The study of animal venoms is of great importance to carry out the characterization of unknown components and that may have a biotechnological application (like the antimicrobial peptides), in special venoms from spiders that are from less studied families.

## Introduction

Several are the animals that evolved using an efficient immune system and excellent protection against pathogens, some of them also employed venoms as a defense mechanism, including arthropods, cnidarian, echinoderms, mollusks, annelids, and vertebrates (Pineda *et al*. 2014, Zhang 2015, Arbuckle 2017).

Concerning biodiversity, spiders are the most successful group of venomous animals (Luddecke *et al*. 2019), comprising more than 50,350 species, 4,280 genera, and 132 families characterized to date (World Spider Catalog 2022). Their venoms are highly elaborate cocktails composed of a heterogeneous mixture of inorganic salts, organic molecules with low molecular mass (< 1 kDa), free amino acids, small polypeptides and linear peptides, nucleotides, disulfide-rich peptides (generally with 3-6 disulfide bonds), as well as enzymes and higher mass proteins (10-120 kDa) (Pineda *et al*. 2014, Saez and Herzig 2019). These molecules can have multiple functions like blockage or modulation of ion channels, anti-cancer, anti-insect, antimicrobial and antihypertensive activities (Melo-Braga *et al*. 2020).

To date, more than 1,390 spider toxins have been described in Uniprot (The UniProt Consortium 2021) most of them from araneomorph spiders, as the anticancer peptide Latarcin-2a from *Lachesana tarabaevi* (Dubovskii *et al*. 2015), or the neurotoxins Tx3, PnPP-19, and δ-CNTX-Pn1a from well-studied *Phoneutria nigriventer* (Cordeiro *et al*. 1992, Cordeiro *et al*. 1993, Figueiredo *et al*. 1995). Although less studied, the suborder Mygalomorphae comprises spiders that are very helpful for biotechnology purposes due to their longevity (15-30 years) and large venom glands (Bond *et al*. 2012, Windley *et al*. 2012). The biggest mygalomorph species are included in the Theraphosidae family and their toxins are named theraphotoxins, whose majority conform to the Inhibitory Cystine Knot (ICK) motif (Escoubas and Rash 2004, King *et al*. 2008). The ICK fold seems to be conserved in several venomous animal lineages and presents elevated functional diversity (Gao *et al*. 2013), as well as gives tremendous thermal, chemical, and biological stability to the toxins (Saez *et al*. 2010).

The peptides GsMTx4, from *Grammostola spatulata* venom (Jung *et al*. 2006), and Oh-defensin, from *Cyriopagopus hainanus* (Zhao *et al*. 2011), are examples of theraphotoxins that adopt the ICK motif and have antimicrobial activity. These toxins are antimicrobial peptides (AMPs), molecules that can be good candidates for the development of new antibiotics since they present notable features like specific action against microorganisms, accelerated pharmacokinetics, immunomodulatory activity, the effect on endotoxin neutralization, and also the possibility of combining with conventional antibiotics to increase the treatment effectiveness (Santos *et al*. 2016).

The search for novel effective antibiotics becomes urgent due to the remarkable proficiency of pathogenic microorganisms at adapting to environmental stresses and developing at least one mechanism of resistance to conventional drugs, compromising their effectiveness. The ever-increasing antibiotic resistance associated with the decrease in the rate of creation of new medicines makes infectious diseases one of the main causes of human deaths (Saez *et al*. 2010, Garcia *et al*. 2013, Saez and Herzig 2019).

Therefore, in this work, we analyzed the venom of the spider *Avicularia juruensis* (Mygalomorphae: Theraphosidae) (Mello Leitão 1922) looking for new antimicrobial peptides. This species is found in Colombia, Ecuador, Peru, and Brazil (Fukushima and Bertani 2017), and is often an exotic pet due to its docility, awesome color, and size. Although *A. juruensis* is easily raised in captivity, there are only two studies about its venom composition, in which it was described the antifungal peptide Juruin (Ayroza *et al*. 2012) and a neprilysin (Nascimento *et al*. 2021).

In addition to the identification of molecules that can be employed in the development of new drugs effective against resistant microorganisms, the study of venoms from mygalomorph spiders is very important to understand its composition, since this suborder is less studied than Araneomorphae.

## Materials and methods

### Animals and venom milking

Spiders from the species *A. juruensis* were collected in Porto Velho and Monte Negro, Rondônia, Brazil, under the Permanent Zoological Material license n°11024-3-IBAMA and Special Authorization for Access to Genetic Patrimony n°010345/2014-0. They were kept in the bioterium of the Laboratory of Applied Toxinology at Butantan Institute (São Paulo, Brazil) and were fed with cockroaches every 15 days, with water at ease. The room temperature was controlled between 24 and 26°C.

Venom was extracted by electrical stimulation as described by Rocha-e-Silva *et al*. (2009). After milking, the venom obtained was centrifuged at 2450× g for 10 minutes at 4°C; the supernatant was collected, lyophilized, and stored at −80°C.

This research follows the Ethical Principles in Animal Research adopted by the Ethic Committee in the Use of Animals of Butantan Institute and was approved on September 21, 2016 (protocol n°CEUAx 5082310816).

### Reverse-phase high-performance liquid chromatography (RP-HPLC)

5 mg of crude venom were reconstituted in 2 mL of acidified ultrapure water [0.05% trifluoroacetic acid (TFA)] and centrifuged at 16,000× g for 5 minutes, the supernatant was subjected to RP-HPLC on a Prominence LC-20A system (Shimadzu, Kyoto, KY, Japan), carried out with a semi-preparative reverse-phase column Jupiter® C18 (10 μm, 300 Å, 10 mm x 250 mm) (Phenomenex International, Torrance, CA, USA).

Two eluents were used: phase A (0.05% TFA) and phase B [acetonitrile (ACN)/0.05% TFA]. Elution was made with a linear gradient of 2 to 80% of phase B over 60 minutes, with a flow rate of 1.5 mL/minute and absorbance monitored at 225 nm. The fractions corresponding to peaks observed in the chromatogram were collected manually and dried in a refrigerated vacuum centrifuge.

### Second chromatography step

The fractions with antimicrobial activity were acidified with 0.05% TFA and subjected to the second chromatography step using the same system aforementioned and an analytic reverse-phase column Jupiter® C18 (10 μm, 300 Å, 250 × 4.60 mm) (Phenomenex). A linear gradient of phase B was calculated specifically for each fraction according to their retention time observed in the first chromatography step: Avilin = from 0 to 7%, Juruin_2 = from 18 to 48%, and Juruenine = from 23 to 53%. Elutions were carried out for 60 minutes, with a flow rate of 1 mL/minute and absorbance monitored at 225 nm. The fractions corresponding to peaks observed in the chromatogram were collected manually and dried in a refrigerated vacuum centrifuge.

### Quantification

Quantification was carried out based on absorbance at 225 nm measured by NanoDrop 2000 spectrophotometer (Thermo Fisher Scientific Inc., Waltham, MA, USA).

### Bioassays

#### Microbial growth inhibition assay and minimum inhibitory concentration (MIC)

Fractions collected in the RP-HPLC step were reconstituted with ultrapure water and tested against Gram-negative bacteria *Escherichia coli* (strain SBS 363), Gram-positive bacteria *Micrococcus luteus* (strain A270), yeast *Candida albicans* (strain MDM8), and the fungi *Aspergillus niger*.

Bacteria were cultured in Poor Broth medium (PB) (1% bactopeptone, 0.5% NaCl, pH 7.4, 217 mOsm) and fungi were cultured in Potato Dextrose Broth medium (PDB) (1.2% PDB, pH 5.0, 79 mOsm). 0.2 mg of streptomycin and sterile ultrapure water were employed as a positive inhibition control and microbial growth control, respectively. The plates were incubated at 30°C for 18 hours under constant shaking, then the antimicrobial activity was evaluated by measuring absorbance at 595 nm in a Victor3 microplate reader (Perkin Elmer Inc., Waltham, MA, USA).

The determination of MIC was performed as described above but using a serial dilution of the antimicrobial fractions. It was expressed as [a]–[b]: [a] is the highest concentration at which the microorganism grows and [b] is the lowest concentration that inhibits the growth.

#### Hemolytic assay

Type A+ human blood was centrifuged at 700× g for 15 min, plasma was discarded and the red blood cells were washed three times in phosphate-buffered saline (PBS) 1X (137 mM NaCl, 2.7 mM KCl, 10 mM Na_2_HPO_4_, 1.76 mM KH_2_PO_4_, pH 7.4), at each washing new centrifugations were performed and the supernatant was discarded. The erythrocytes obtained were diluted in the same PBS 1X buffer until a final concentration of 3% (v/v).

Aliquots of the antimicrobial fractions were reconstituted in the PBS 1X buffer and tested in duplicate in serial dilutions using a 96-well U-bottom microplate. 0.1% Triton X-100 solution and the PBS 1X buffer were used as positive and negative hemolysis control, respectively. The plate was incubated for 1 hour at 37°C, then the supernatant was transferred to a 96-well flat-bottom microplate, and hemolysis was evaluated by measuring the absorbance at 405 nm using a Victor3 microplate reader (Perkin Elmer). Hemolysis percentage was expressed concerning the 100% lysis control, according to the equation: % hemolysis = (Absorbance sample - Absorbance PBS 1X)/(Absorbance Triton X-100 - Absorbance PBS 1X).

The use of human blood was authorized by the Brazilian National Health Council, Plataforma Brasil CAAE: 15390119.9.0000.5473 (protocol number 3.486.142).

#### Cytotoxicity assay

Vero cells (ATCC CCL81; Manassas, VA, USA) were cultivated and maintained in Opti-MEM(1X) + GlutaMAX (reduced serum medium) culture medium (Gibco, Waltham, MA, USA). The cells were seeded in 96-well flat-bottom sterile plates (2 × 10^5^ cells/well) and cultured for 24 h at 37°C in a humidified atmosphere containing 5% CO_2_. Serial dilutions of the samples were carried out using the same culture medium aforementioned (starting at 100 μg/mL for Avilin and 100 μM for Juruin_2 and Juruenine) and were incubated with cells for 24 h. The assays were conducted in duplicate and the culture medium was employed as a negative cytotoxicity control. 20 μL of MTT (5 mg/mL in PBS) were added and a new incubation was carried out for 3 h at 37°C. After, 150 μL of isopropanol was used to dissolve the formazan crystals.

Absorbance at 570 nm was measured using a FlexStation 3 microplate reader (Molecular Devices, Sunnyvale, CA, USA) and the cell survival was calculated according to the equation: % survival = (Absorbance treated cells/Absorbance untreated cells) × 100.

### Transcriptomic analysis and database assembly

Venom glands of five adult spiders were removed and homogenized in Polytron® Tissue Homogenizer. Total RNA was extracted using TRIzol™ Reagent (Thermo Fisher Scientific Inc., Waltham, MA, USA) and mRNA was isolated with magnetic beads with an oligo (dT) using a Dynabeads® mRNA DIRECT™ kit (Thermo). It was quantified using Quant-iT™ RiboGreen® RNA Assay Kit (Thermo) and its integrity was evaluated in a 2100 Bioanalyzer pico chip series (Agilent Technologies, Santa Clara, CA, USA).

The mRNA obtained was used to assemble the cDNA library, following the standard TruSeq RNA Sample Prep Kit protocol (Illumina, San Diego, CA, USA). The cDNA library was sequenced on Illumina HiSeq 1500 system, into a rapid paired-end flow cell in 300 cycles of 2*150 bp paired-end technique, according to the standard manufacturer’s protocol (Illumina). Transcripts obtained were assembled using rnaSPAdes software, version 3.10.1 (Bushmanova *et al*. 2019), and filtered according to the minimum size of 300 bp. Also, redundant transcripts were removed using CD-HIT software (Li and Godzik 2006, Fu *et al*. 2012).

Potential coding sequences were identified by TransDecoder software, version 2.0.1 (https://github.com/TransDecoder/TransDecoder/wiki/), with a minimum protein length of 60. Transcripts containing the translated sequences were aligned by BLASTp (Altschul *et al*. 1990) against the databases UniProt/Swiss-Prot proteins and TSA-NR from NCBI to assess the protein description with a cut-off e-value of <1e-05 and according to the criterion with longer protein similarity. The predicted proteins were analyzed for Protein families (Pfam) domains with HMMER3.1 (Wheeler and Eddy 2013), against the Pfam domains database (El-Gebali *et al*. 2019). TransDecoder may predict more than one coding sequence by transcript and we selected only the best one based on the priority order of UniProt-KB/Swiss-Prot, Pfam domains database, and TSA-NR. All these databases were used for annotation and selection of the best candidate for each transcript.

### In-solution digestion (reduction, alkylation, and trypsinization)

Aliquots of antimicrobial fractions were reconstituted in 20 μL of 8 M urea/0.4 M ammonium bicarbonate solution. 5 μL of reducing solution [45 mM dithiothreitol (Invitrogen, Carlsbad, CA, USA)] were added and incubated at 50°C for 15 minutes. After cooling at room temperature, 5 μL of alkylation solution [100 mM iodoacetamide (GE Healthcare, Chicago, IL, USA)] was added and incubated protected from light for 15 minutes. 130 μL of ultrapure water and bovine trypsin (Sigma-Aldrich, San Luis, MO, USA) were added in a 1:25 (enzyme:protein) ratio and incubated at 37°C for 12 hours, then the digestion was stopped by acidifying with 0.1% TFA.

The samples were desalinated by ZipTip® Pipette Tips (Merck Millipore, Billerica, MA, USA) with a unique elution step using 80% ACN, concentrated in a vacuum centrifuge, and analyzed by mass spectrometry.

### Mass spectrometry and bioinformatics analyses

Aliquots of the antimicrobial fractions (trypsinized and non-trypsinized) were reconstituted in 0.1% formic acid solution and analyzed by the Easy-nLCII system (Thermo) connected to LTQ-Orbitrap Velos (Thermo). Electrospray ionization was operated in positive mode, with voltage and temperature adjusted to 2.0 kV and 200°C. The mass scan interval considered for the full scan (MS1) was 200-2,000 m/z, operating in data-dependent acquisition mode. The five most intense ions were selected for collision-induced dissociation fragmentation and the minimum signal required to start fragmentation events (MS2) was adjusted to 5,000 cps.

The MS/MS spectra obtained from non-trypsinized samples were analyzed by MassAnalyzer software, version 1.03 (Amgen, Thousand Oaks, CA, USA). The spectra of trypsinized samples were converted to a .mgf file by MSConvert software (Chambers *et al*. 2012) and employed in searches in different databases using Mascot Server (Perkins *et al*. 1999). They were also processed by PEAKS Studio software, version X Plus (Bioinformatics Solution Inc., Waterloo, ON, Canada). Using the PEAKS DB function, the spectra were searched against the database assembled with predicted proteins obtained in the transcriptomic analysis. The searches involved 10 ppm and 0.6 Da error tolerance for precursor ions and fragment ions, respectively.

Database searches were performed using BLASTp and Uniprot. Alignments and coloring were made using Seaview (Gouy *et al*. 2010) and Sequence Manipulation Suite (Stothard 2000), respectively. The similarity percentage was calculated by the SIAS server (http://imed.med.ucm.es/Tools/sias.html), and physical and chemical parameters were evaluated by ProtParam (Gasteiger *et al*. 2005). For all tools were used default parameters.

## Results

### Fractionation of the venom by RP-HPLC

To isolate and characterize the antimicrobial peptides (AMPs), *A. juruensis* venom was fractionated by RP-HPLC, and all fractions obtained were tested by microbial growth inhibition assay. Thus, were identified three fractions with antimicrobial activity: fraction 1, which prevented the growth of *Escherichia coli* SBS 363; fraction 19, which prevented the growth of *Aspergillus niger, Candida albicans* MDM8, *Micrococcus luteus* A270, and *E. coli* SBS 363; and fraction 24, which was effective against *E. coli* SBS 363 and *M. luteus* A270.

By mass spectrometry analysis, it was observed that these fractions were not homogeneous (data not shown), therefore, it was necessary to submit them to a new RP-HPLC step. The purified antimicrobial fractions obtained were named Avilin, Juruin_2, and Juruenine (Figure 1 and Appendices 1 to 3).

**Figure 1.**
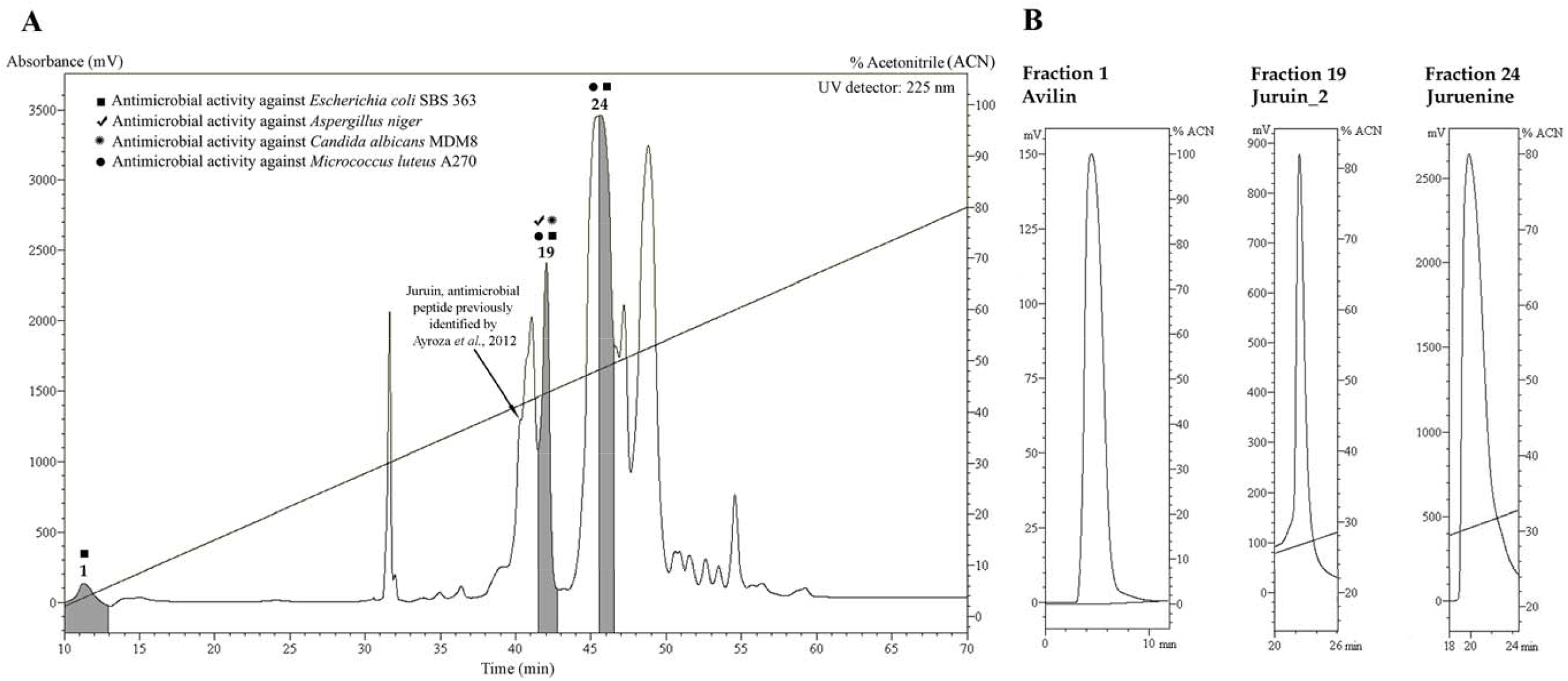
(A) Chromatographic profile of *Avicularia juruensis* venom, obtained by high-performance liquid chromatography carried out with a semi-preparative reverse-phase column Jupiter® C18 (Phenomenex). Elution was performed using a gradient of 2 to 80% of acetonitrile in acidified water, over 60 minutes under a flow rate of 1.5 mL/minute. The fractions corresponding to the peaks observed in the chromatogram were collected separately, those with antimicrobial activity are highlighted: fraction 1, which prevented the growth of the Gram-negative bacteria *Escherichia coli* SBS 363; fraction 19, which prevented the development of the fungus *Aspergillus niger*, the yeast *Candida albicans* MDM8, the Gram-positive bacteria *Micrococcus luteus* A270 and *E. coli* SBS 363; and fraction 24, which prevented the growth of *E. coli* SBS 363 and *M. luteus* A270. The fraction corresponding to the antimicrobial peptide Juruin (previously identified in *A. juruensis* venom by Ayroza *et al*., 2012) is also indicated. (B) Chromatographic profiles of the second high-performance liquid chromatography step performed with the fractions 1, 19, and 24. Elutions were carried out using an analytic reverse-phase column Jupiter® C18 (Phenomenex) under a flow rate of 1 mL/minute for 60 minutes. A linear gradient of acetonitrile in acidified water was calculated specifically for each sample according to their retention time observed in the first chromatography step: fraction 1 = from 0 to 7%, fraction 19 = from 18 to 48%, and fraction 24 = from 23 to 53%). The purified antimicrobial fractions obtained were named Avilin, Juruin_2, and Juruenine.

### Bioassays

The Table 1 indicates the MIC of Avilin, Juruin_2, and Juruenine against the microorganisms tested. By hemolytic assay, it was observed that only Avilin has toxic effects in MIC concentrations (Figures 2C, 3D, and 4D). Concerning cytotoxicity assay, the viability of cells decreased at the maximum concentrations tested for all samples, and Juruenine is the molecule with the lowest cytotoxicity observed (cell viability between 70-90%) (Figures 2D, 3E, and 4E).

**Table 1.** Minimum inhibitory concentration (MIC) of antimicrobial fractions.

| Microorganism | Avilin |  | Juruin_2 |  | Juruenine |  |
| --- | --- | --- | --- | --- | --- | --- |
| | MIC<br>(μM) | MIC<br>(μg/mL) | MIC<br>(μM) | MIC<br>(μg/mL) | MIC<br>(μM) | MIC<br>(μg/mL) |
| *Aspergillus niger* | - | - | 3.2 - 6.4 | 13.8 - 27.6 | - | - |
| *Candida albicans* MDM8 | - | - | 1.6 - 3.2 | 6.9 - 13.8 | - | - |
| *Escherichia coli* SBS 363 | ND | 1.75 - 3.5 | 1.6 - 3.2 | 6.9 - 13.8 | 2.5 - 5 | 10.46 - 20.92 |
| *Micrococcus luteus* A270 | - | - | 1.6 - 3.2 | 6.9 - 13.8 | 20 - 40 | 83.67 - 167.35 |

Avilin, Juruin\_2, and Juruenine were tested against the fungi *Aspergillus niger*, the yeast *Candida albicans* (strain MDM8), the Gram-negative bacteria *Escherichia coli* (strain SBS 363), and the Gram-positive bacteria *Micrococcus luteus* (strain A270). Symbol (-): not detected. ND: not determinate.

**Figure 2.**
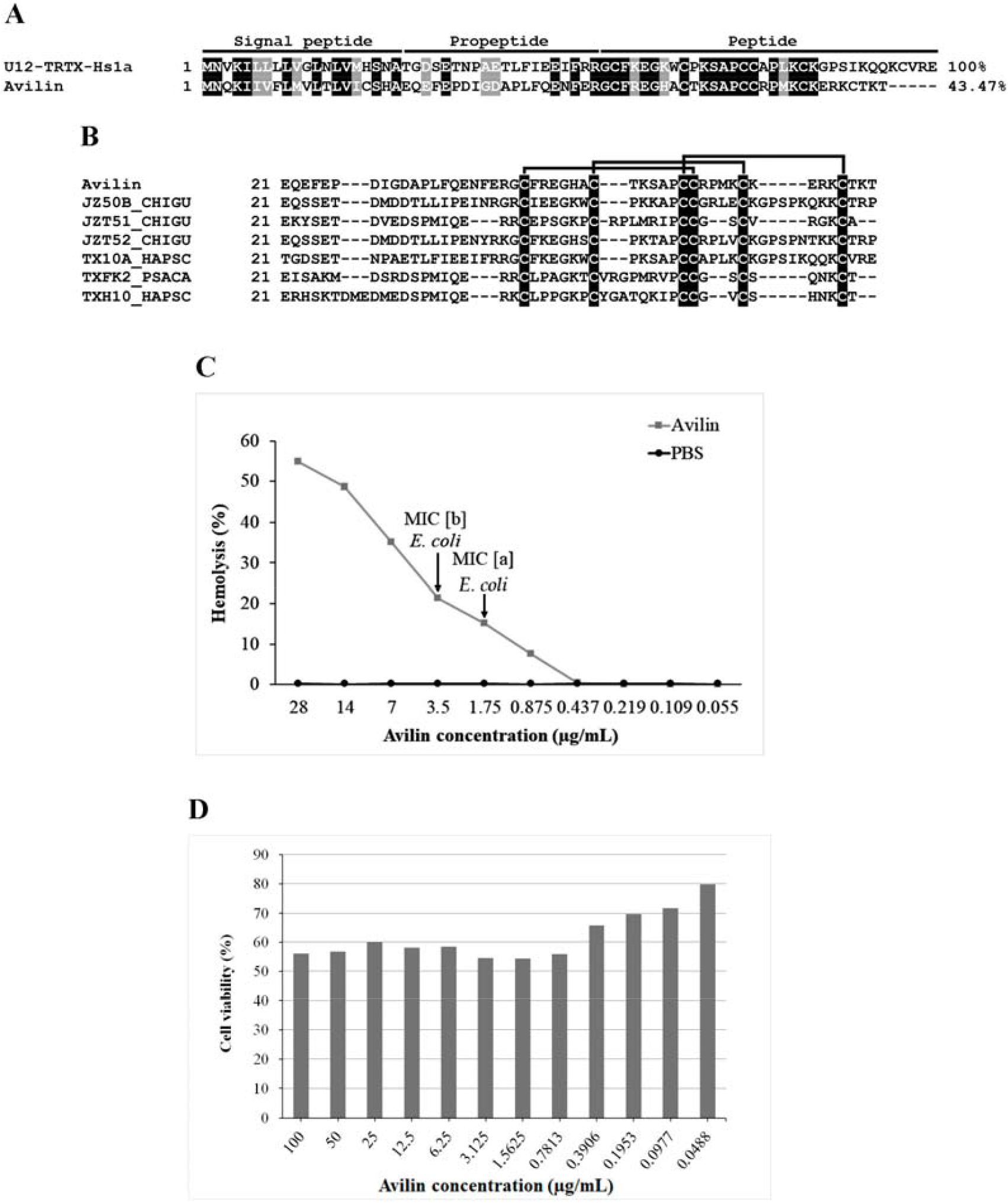
(A) Alignment of Avilin with the theraphotoxin U12-TRTX-Hs1a from the venom of the spider *Cyriopagopus schmidti*. The signal peptide, propeptide, and active peptide are indicated. Conserved amino acids are shown in black, those with the same chemical character are shown in gray, and non-conserved amino acids are shown in white. The percentages of identity, compared to the first sequence, are indicated at the end of the amino acid sequence. The symbol (-) represents the gaps. (B) Alignment of Avilin with five theraphotoxins that have the Inhibitor Cystine Knot (ICK) motif. The disulfide bridges are formed between C1-C4, C2-C5, and C3-C6 (highlighted in black). Uniprot ID JZT52_CHIGU: theraphotoxin U5-TRTX-Cg1a from the venom of the spider *Chilobrachys guangxiensis*. TX10A_CYRSC: theraphotoxin U12-TRTX-Hs1a from *Cyriopagopus schmidt*. JZ50B_CHIGU, JZ50A_CHIGU, and JZ50C_CHIGU: isoforms of the theraphotoxin U4-TRTX-Cg1a from *C. guangxiensis*. (C) Hemolytic effect of Avilin on human erythrocytes. The concentration-response curves of hemolytic activity show its toxicity at different concentrations. The arrows indicate the minimum inhibitory concentration at which Avilin has antimicrobial activity against *Escherichia coli* SBS 363 ([a] is the highest concentration at which the microorganism grows and [b] is the lowest concentration that inhibits the growth). PBS was used as a negative hemolysis control. (D) Cytotoxic effect of Avilin on Vero cells (ATCC CCL81). The cells were incubated with different concentrations of Avilin (0.05 to 100 μg/mL) for 24 h at 37°C. The effects on cell viability were determined by MTT colorimetric assay. Untreated cells were used as a negative cytotoxic control.

**Figure 3.**
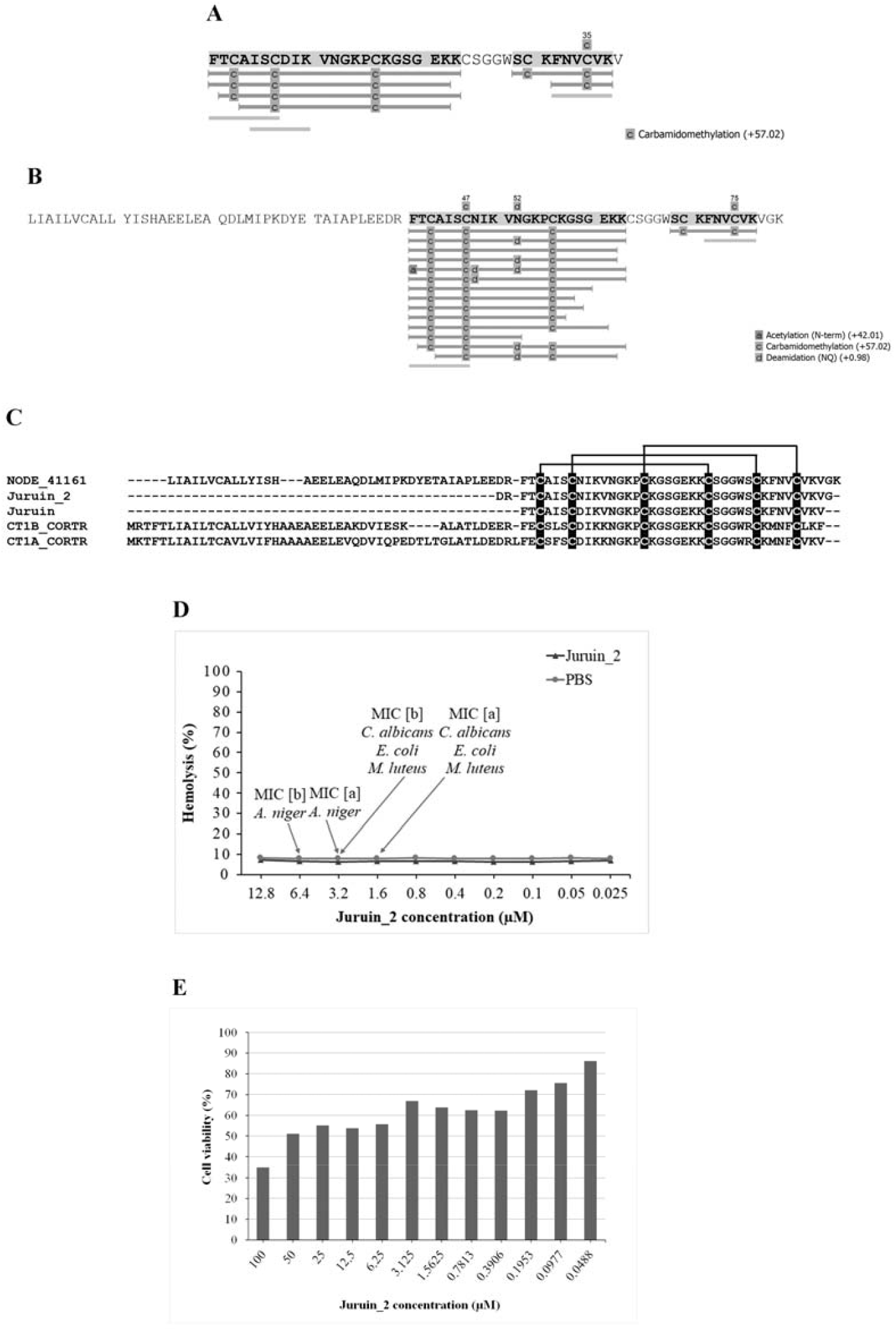
(A) Result obtained in mass spectrometry analyses of Juruin_2, carried out by PEAKS Studio software X Plus (Bioinformatics Solution). Juruin_2 has peptide fragments corresponding to the antimicrobial peptide Juruin (Ayroza *et al*., 2012), which are highlighted by gray shadings and bars. The carbamidomethylation of cysteines (post-translational modification) is indicated. (B) By mass spectrometry, it was also possible to identify that the NODE_41161 encodes the Juruin and Juruin_2. The peptide fragments detected are highlighted by gray shadings and bars. N-term acetylation, carbamidomethylation of cysteines, and NQ deamidation (post-translational modification) are indicated. (C) Alignment of the translated amino acids of NODE_41161, Juruin, and Juruin_2 with two theraphotoxins that have the Inhibitor Cystine Knot (ICK) motif. The disulfide bridges are formed between C1-C4, C2-C5, and C3-C6 (highlighted in black). Uniprot ID CT1B_CORTR: theraphotoxin U1-TRTX-Ct1b from the venom of the spider *Coremiocnemis tropix*. CT1A_CORTR: theraphotoxin U1-TRTX-Ct1a from *C. tropix*. (D) Hemolytic effect of Juruin_2 on human erythrocytes. The concentration-response curves of hemolytic activity show its toxicity at different concentrations. The arrows indicate the minimum inhibitory concentration at which Juruin_2 has antimicrobial activity against *Escherichia coli* SBS 363, *Micrococcus luteus* A270, *Candida albicans* MDM8, and *Aspergillus niger*. ([a] is the highest concentration at which the microorganism grows and [b] is the lowest concentration that inhibits the growth). PBS was used as a negative hemolysis control. (E) Cytotoxic effect of Juruin_2 on Vero cells (ATCC CCL81). The cells were incubated with different concentrations of Juruin_2 (100 to 0.0488 μM) for 24 h at 37°C. The effects on cell viability were determined by MTT colorimetric assay. Untreated cells were used as a negative cytotoxic control.

**Figure 4.**
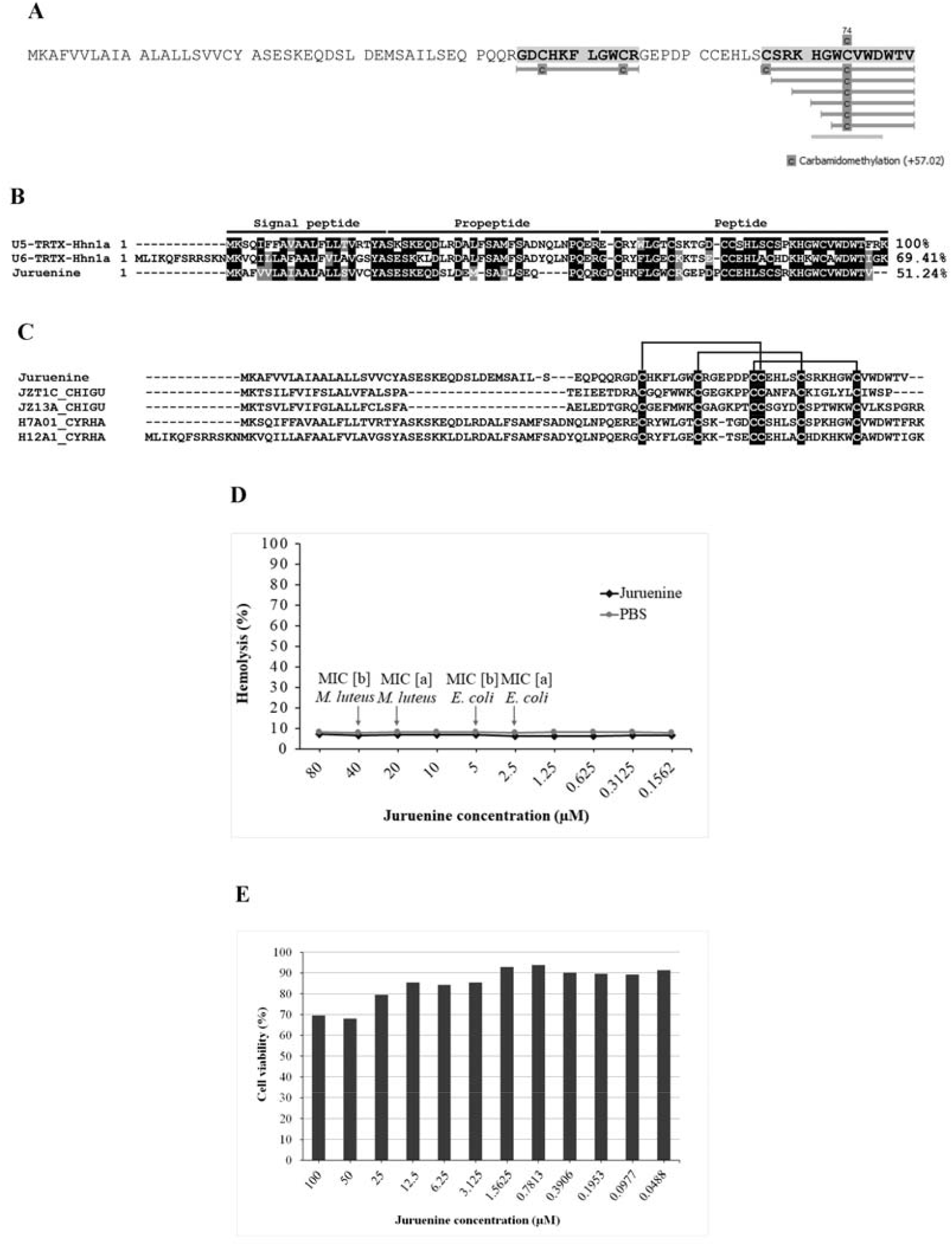
(A) Result obtained in mass spectrometry analyses of Juruenine, carried out by PEAKS Studio software X Plus (Bioinformatics Solution). Juruenine has peptide fragments corresponding to the NODE_17940, which are highlighted by gray shadings and bars. The carbamidomethylation of cysteines (post-translational modification) is indicated. (B) Alignment of Juruenine with the theraphotoxins U5-TRTX-Hhn1a and U6-TRTX-Hhn1a from the venom of the spider *Haplopelma hainanum*. The signal peptide, propeptide, and active peptide are indicated. Conserved amino acids are shown in black, those with the same chemical character are shown in gray, and non-conserved amino acids are shown in white. The percentages of identity, compared to the first sequence, are indicated at the end of the amino acid sequence. The symbol (-) represents the gaps. (C) Alignment of Juruenine with four theraphotoxins that have the Inhibitor Cystine Knot (ICK) motif. The disulfide bridges are formed between C1-C4, C2-C5, and C3-C6 (highlighted in black). Uniprot ID JZT1C_CHIGU: theraphotoxin Delta-TRTX-Cg1a from the venom of the spider *Chilobrachys guangxiensi*. JZ13A_CHIGU: theraphotoxin U10-TRTX-Cg1a from *Coremiocnemis tropix*. H7A01_CYRHA: theraphotoxin U5-TRTX-Hhn1a from *Cyriopagopus hainanus*. H12A1_CYRHA: theraphotoxin U6-TRTX-Hhn1a from *C. hainanus*. (D) Hemolytic effect of Juruenine on human erythrocytes. The concentration-response curves of hemolytic activity show its toxicity at different concentrations. The arrows indicate the minimum inhibitory concentration at which Juruenine has antimicrobial activity against *Escherichia coli* SBS 363 and *Micrococcus luteus* A270 ([a] is the highest concentration at which the microorganism grows and [b] is the lowest concentration that inhibits the growth). PBS was used as a negative hemolysis control. (E) Cytotoxic effect of Juruenine on Vero cells (ATCC CCL81). The cells were incubated with different concentrations of Juruenine (100 to 0.0488 μM) for 24 h at 37°C. The effects on cell viability were determined by MTT colorimetric assay. Untreated cells were used as a negative cytotoxic control.

### Characterization of antimicrobial fractions

#### Avilin (Fraction 1)

No significant results were obtained in the analyzes carried out with PEAKS X and MassAnalyzer. In the searches performed with the Mascot Server, it was observed that Avilin has a similarity with the theraphotoxin U12-TRTX-Hs1a from the venom of the spider *Cyriopagopus schmidti*. Using this theraphotoxin as a reference were performed searches among the predicted proteins obtained in the transcriptomic analysis, thus, it was identified the transcript which encodes the Avilin (NODE_18923), as well as their complete sequence with signal peptide, propeptide, and active peptide (Figure 2A and Appendice 4).

By aligning Avilin with five other theraphotoxins which has ICK motif, it is observed that it also probably has this structure, with disulfide bridges formed between C1-C4, C2-C5, and C3-C6 (Figure 2B). Additionally, using ProtParam the molecular mass of the active peptide region of Avilin was evaluated at 3,260.9 Da. Its cationic character was also verified due to the presence of eight positively charged amino acids (five lysines and three arginines), with pI calculated at 9.08.

#### Juruin_2 (Fraction 19)

By analyzes carried out with PEAKS X, it was observed high similarity with the antimicrobial peptide Juruin, which was previously isolated and characterized in *A. juruensis* venom by Ayroza et al. (2012) (Figure 3A). Additionally, it was identified that the transcript NODE_41161 encodes the Juruin and Juruin_2 (Figure 3B and Appendice 5).

Analyses carried out with Mascot Server indicated that Juruin_2 has similarity with the theraphotoxin U4-TRTX-Hhn1ac from the venom of the spider *Cyriopagopus hainanus* and, as expected, it has ICK motif with disulfide bridges formed between C1-C4, C2-C5, and C3-C6 (Figure 3C).

Searches performed using MassAnalyzer indicated a molecular mass of 4,319.1 Da (Appendice 6), which corresponds to the Juruin molecule with DR and G amino acids linked to its N and C-terminal portions, respectively (**DR**FTCAISCNIKVNGKPCKGSGEKKCSGGWSCKFNVCVKV**G**). This result shows that even with three more amino acid residues linked to its original structure, Juruin still maintains its antimicrobial action, and this isoform was named Juruin_2.

By ProtParam was verified that Juruin_2 has a cationic character due to the presence of eight positively charged amino acids (seven lysines and one arginine), with pI calculated at 9.27.

#### Juruenine (Fraction 24)

No significant results were obtained in the analyzes carried out with the Mascot Server. In the searches performed with PEAKS X, it was observed that Juruenine contains peptide fragments corresponding to the transcript NODE_17940 (Figure 4A and Appendice 7). Using BLASTp it was identified similarity with the theraphotoxins U5-TRTX-Hhn1a and U6-TRTX-Hhn1a from the venom of the spider *Cyriopagopus hainanus*. By alignment and comparison with these toxins, the signal peptide, propeptide, and active peptide were identified (Figure 4B).

As Avilin and Juruin_2, Juruenine probably has an ICK motif in this structure, with disulfide bridges formed between C1-C4, C2-C5, and C3-C6 (Figure 4C). Searches performed using MassAnalyzer indicated a molecular mass of 4,221.6 Da (Appendice 8), which corresponds to the fragment GDCHKFLGWCRGEPDPCCEHLSCSRKHGWCVWDWTV, the active peptide of Juruenine. Additionally, by ProtParam it was verified that the active peptide has an anionic character due to the presence of eleven negatively charged amino acids (five aspartic acid and six glutamic acid), with pI calculated at 5.46.

## Discussion

The study of antimicrobial molecules is an important strategy to identify substances that may be alternatives to actual antibiotics. Several works carried out with invertebrates characterized antimicrobial peptides from the hemolymph of *Acanthoscurria rondoniae* (Riciluca *et al*. 2012), *Acutisoma longipes* (Sayegh *et al*. 2016), *Tityus serrulatus* (de Jesus Oliveira *et al*. 2019), and *Triatoma infestans* (Diniz *et al*. 2020), for example. Concerning spider venoms, it was verified the presence of these molecules in the species *Acanthoscurria gomesiana* (Abreu *et al*. 2017), *Lycosa erythrognatha* (Santos *et al*. 2010), *L. singoriensis* (Wang *et al*. 2016), and *Lachesana tarabaevi* (Lazarev *et al*. 2013), for example.

All antimicrobial molecules identified in *A. juruensis* venom are effective against *E. coli* SBS 363 and Avilin has the lowest MIC (1.75 - 3.5 μg/mL), nether than the MIC of the peptide U1-SCRTX-Lg1a from *Loxosceles gaucho* venom (7.6 μg/mL) (Segura-Ramírez and Silva Júnior 2018). Only Juruin_2 prevented the growth of all microorganisms tested, similar to Gomesin, a potent antimicrobial peptide isolated from hemocytes of mygalomorph spider *A. gomesiana* (Silva Júnior *et al*. 2000). However, this peptide has MIC up to four times smaller than Juruin_2. Juruenine showed antimicrobial activity against *M. luteus* A270 and *E. coli* SBS 363, but with a MIC eight times higher for Gram-positive bacteria than Gram-negative, probably due to differences between the cell walls of these bacterias.

In the hemolytic assays, it was observed only Avilin causes hemolysis of human erythrocytes from a concentration of 0.875 μg/mL. This characteristic is also observed for the Zodatoxins Ltc 1, Ltc 2a, and Ltc 5 from *L. tarabaevi* venom (Kozlov *et al*. 2006) and Lycosin-II from *L. singoriensis* venom (Wang *et al*. 2016). The Zodatoxins cause hemolysis of rabbit red blood cells at concentrations ranging from 245.6 to 17.4 μg/mL, values much higher than those observed for Avilin. The Lycosin-II led to approximately 20% hemolysis of human red blood cells from the concentration of 50 μM.

In cytotoxicity assays, Juruin_2 was the most toxic molecule. However, this cytotoxicity is observed at higher concentrations of which there is antimicrobial activity. The highest MIC is 6.4 μM for *A. niger* and the cytotoxicity is detected from the concentration of 12.5 μM. Furthermore, it was also observed that Juruin_2 is an isoform of Juruin, previously isolated in *A. juruensis* venom by Ayroza *et al*. (2012). In the transcriptomic analysis of the venom glands, the complete nucleotide sequence of the transcript NODE_41161 was not obtained, which made it impossible to identify the signal peptide and propeptide of the Juruin and Juruin_2 molecule. However, it was possible to observe that the amino acid located at position 8 is aspartic acid and not asparagine as described by Ayroza *et al*., (2012). This can be explained by the fact that the molecular masses of these two amino acids are very similar: aspartic acid has a mass of 115 Da, while asparagine has a mass of 114 Da. Moreover, in the study carried out by Ayroza *et al*., (2012) only the antifungal activity of Juruin was observed, while in this work it was detected the antimicrobial activity of Juruin_2 was also against Gram-positive and Gram-negative bacteria.

Avilin has a similarity with the theraphotoxin U12-TRTX-Hs1a from the venom of the spider *Cyriopagopus schmidti*, a neurotoxin that acts by blocking N-type calcium channels (Jiang *et al*. 2008). Juruin_2 and Juruenine have a similarity with the theraphotoxins U4-TRTX-Hhn1ac, U5-TRTX-Hhn1a, and U6-TRTX-Hhn1a from *C. hainanus* (Tang *et al*. 2010), respectively. All these theraphotoxins are also neurotoxins that act in ion channels. According to Redaelli *et al*. (2010), arachnid neurotoxins can also be potent antimicrobial molecules as, for example, the Cupienins 1a, 1b, 1c, and 1d from *Cupiennius salei* venom (Kuhn-Nentwig *et al*. 2002), the Lycotoxins M-LCTX-Hc1a, M-LCTX-Hc2a, and M-LCTX-Ls2a from *Lycosa carolinensis* (Yan and Adams 1998), CsTx-1 from *Cupiennius salei* (Kuhn-Nentwig *et al*. 2012), and GsMTx-4 (Jung *et al*. 2006) from *Grammostola spatulata*.

Avilin, Juruin_2, and Juruenine probably have the ICK motif in their structure. According to Fujitani *et al*. (2007), the ICK motif is fundamental for the antimicrobial activity of Takhistatins, antimicrobial peptides isolated from the horseshoe crab *Tachypleus tridentatus*. The authors observed that Tachystatins A and B are molecules with high similarity, however, in the antimicrobial tests performed, it was noticed that only Tachystatin A is effective against *E. coli*. The difference between the two molecules is in the size of the loop formed between two cysteine residues that form a disulfide bridge. Takhistatin B has a smaller loop, which impairs its antimicrobial potential. Therefore, the ICK motif may also be of fundamental importance for the structure and function of spider venom antimicrobial molecules.

The phylogeny of toxins with ICK motif suggests that they are derived from genes related to β-defensins, a class of innate immunity-related antimicrobial peptides widely found in different living organisms. These toxins may have originated by duplication of genes related to defensins in venom glands, developing new functions during evolution (Fry *et al*. 2009). The existence of β-defensins from *Drosophila melanogaster* reinforces the aforementioned hypothesis. These antimicrobial peptides act by blocking sodium channels, identical to the mechanism of action of the majority of neurotoxins from arachnids that have an ICK motif (Cohen *et al*. 2009).

## Conclusion

In summary, we analyzed the venom of the mygalomorph spider *Avicularia juruensis* and identified 3 new antimicrobial peptides, since they have a molecular mass of less than 10 kDa. Antimicrobial peptides are fundamental molecules from innate immunity found in all living organisms and they are a principal first line of defense against pathogens. Their presence in venom is very important to protect venom glands and, furthermore, considering that the envenomation is the first step of feeding, they could avoid the contamination of the prey by pathogenic microorganisms.

The study of animal venoms is of great importance to carry out the characterization of unknown components and that may have a biotechnological application, in special of spider venoms that are from less studied families compared to those with medical importance.

## Acknowledgments

We would like to thank MSc Thiago de Jesus Oliveira for his help in the maintenance of the bioterium where the spiders used in this study were kept. Mariana Salgado Loureiro de Caldas Morone, Ursula Castro de Oliveira, and Milton Yutaka Nishiyama-Jr for their support in the transcriptomic analysis. Ismael Feitosa Lima for his support in the mass spectrometry analysis.

## Declaration of interest statement

No potential conflict of interest was reported by the authors.

## Funding

This study was financed in part by the Coordenação de Aperfeiçoamento de Pessoal de Nível Superior - Brasil (CAPES) - Finance Code 001, as well as by the Research Support Foundation of the State of São Paulo (FAPESP/CeTICS - grant No. 2013/07467-1) and the Brazilian National Council for Scientific and Technological Development (CNPq - grant No. 472744/2012-7).

## Appendices

**Appendice 1:**
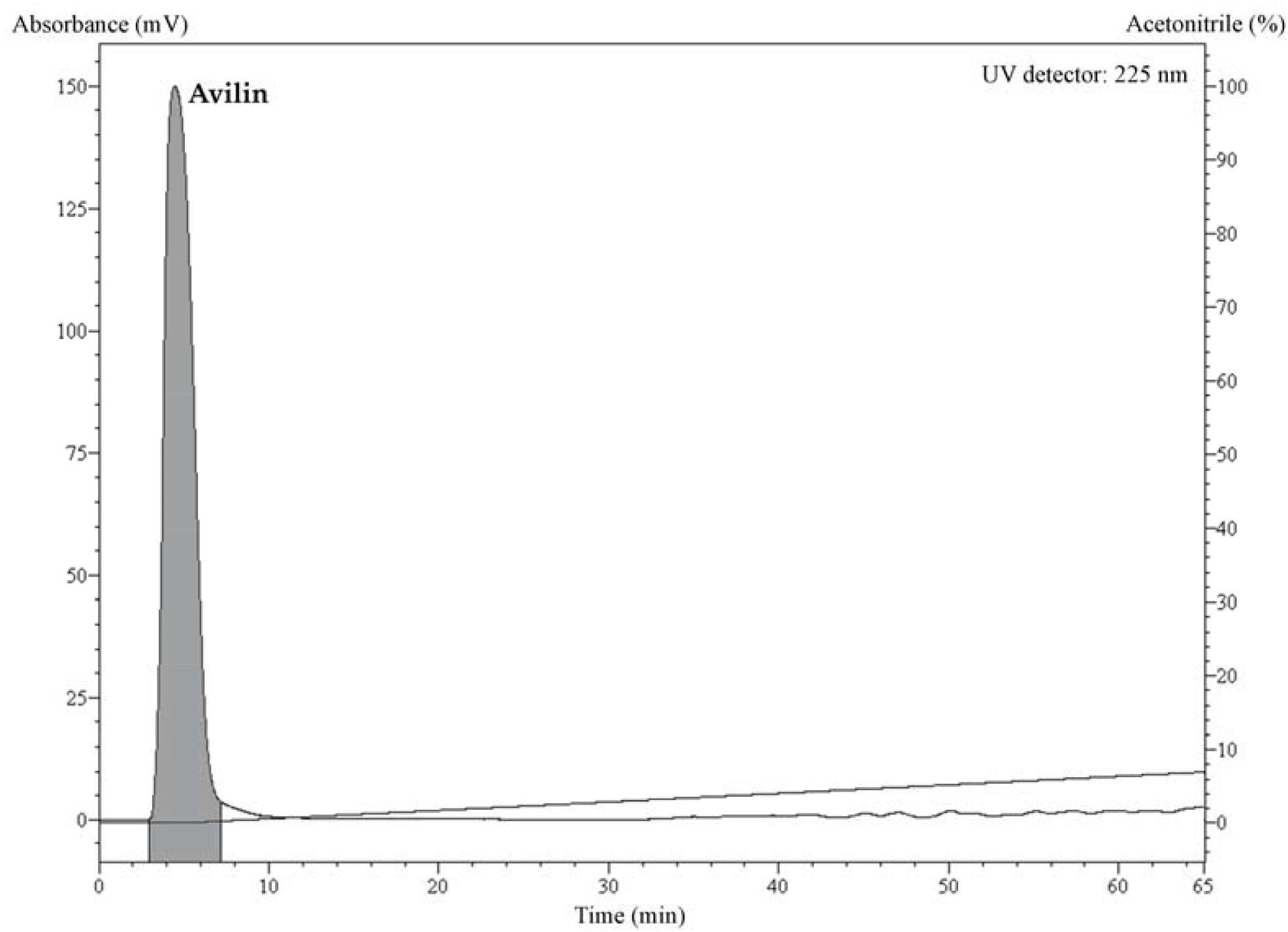
Chromatographic profile of the second high-performance liquid chromatography step performed with the fraction 1. Elution was carried out using an analytic reverse-phase column Jupiter® C18 (Phenomenex) under a flow rate of 1 mL/minute for 60 minutes. A linear gradient of acetonitrile in acidified water was calculated according to its retention time observed in the first chromatography step (from 0 to 7%). The purified antimicrobial fraction obtained was named Avilin.

**Appendice 2:**
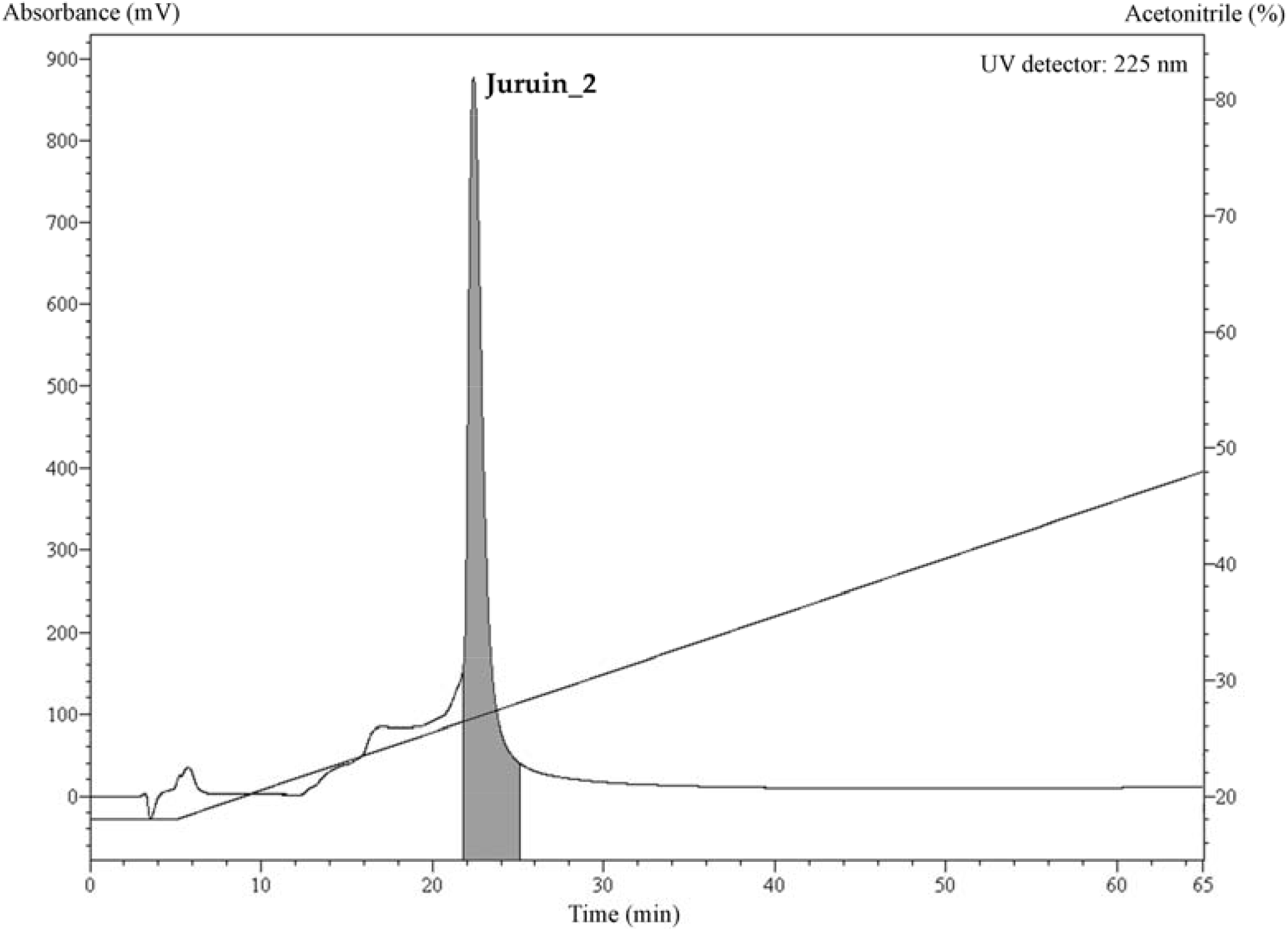
Chromatographic profile of the second high-performance liquid chromatography step performed with the fraction 19. Elution was carried out using an analytic reverse-phase column Jupiter® C18 (Phenomenex) under a flow rate of 1 mL/minute for 60 minutes. A linear gradient of acetonitrile in acidified water was calculated according to its retention time observed in the first chromatography step (from 18 to 48%). The purified antimicrobial fraction obtained was named Juruin_2.

**Appendice 3:**
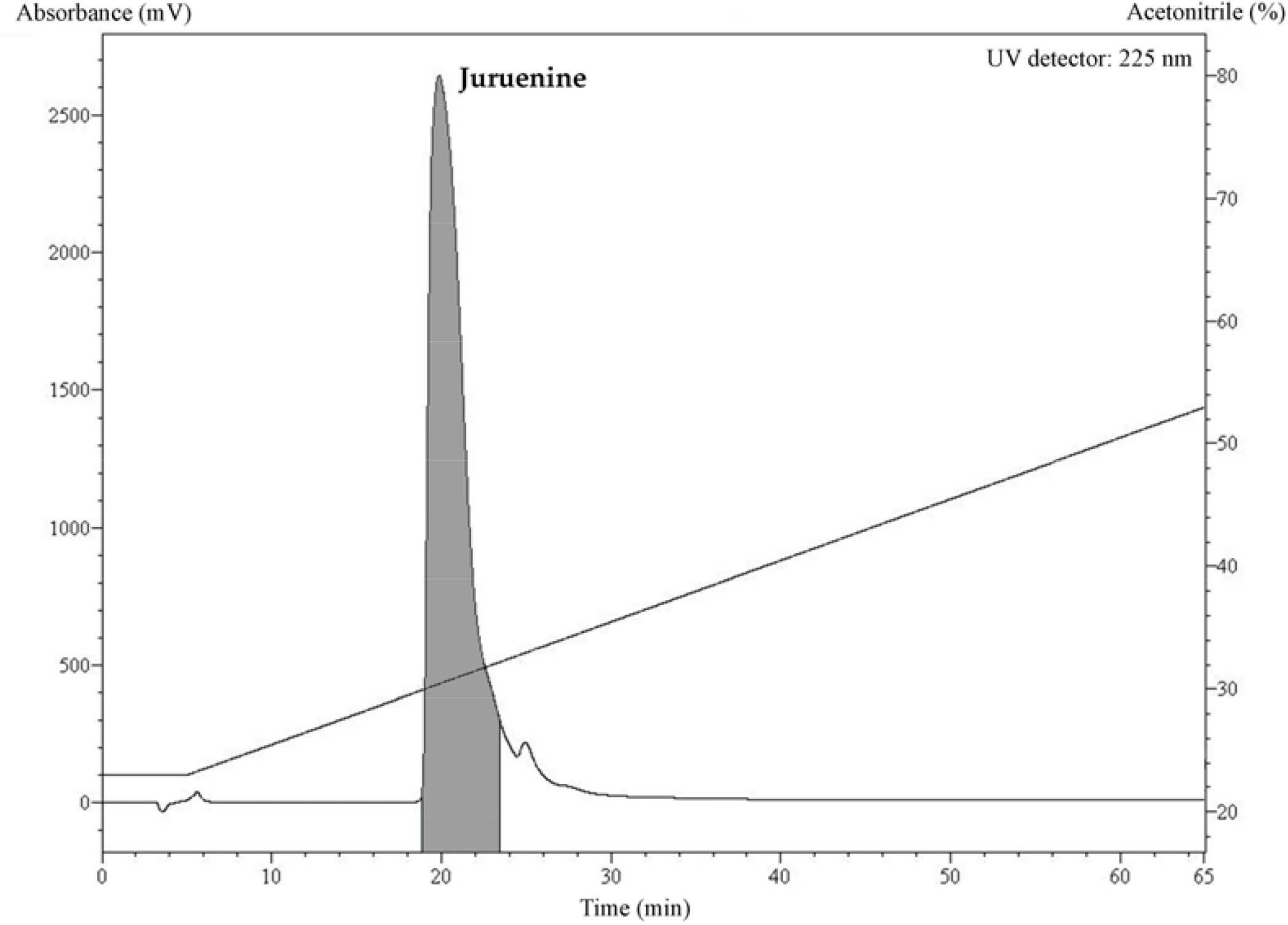
Chromatographic profile of the second high-performance liquid chromatography step performed with the fraction 24. Elution was carried out using an analytic reverse-phase column Jupiter® C18 (Phenomenex) under a flow rate of 1 mL/minute for 60 minutes. A linear gradient of acetonitrile in acidified water was calculated according to its retention time observed in the first chromatography step (from 23 to 53%). The purified antimicrobial fraction obtained was named Juruenine.

**Appendice 4:**
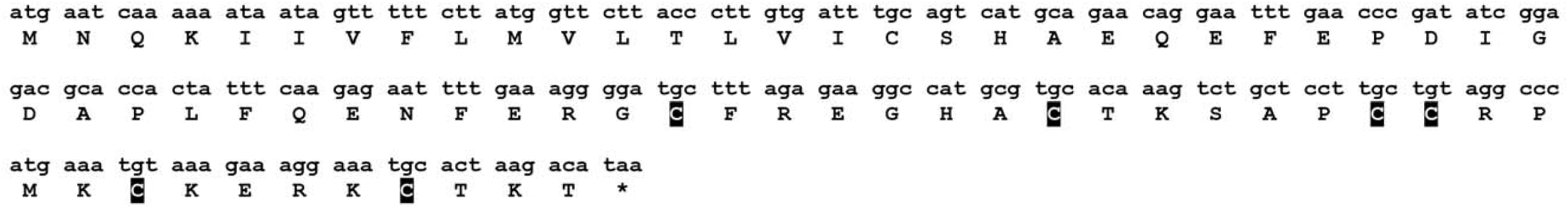
Nucleotide sequence and translated amino acids of NODE_18923 (Xpression_FPKM 943.26), which encodes the Avilin. The cysteines that form disulfide bonds are highlighted in black.

**Appendice 5:**
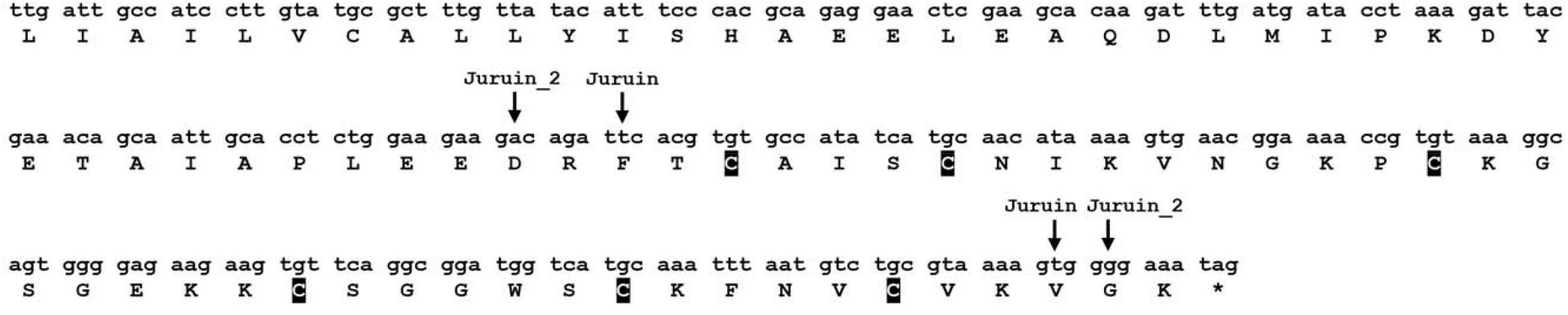
Nucleotide sequence and translated amino acids of NODE_41161 (Xpression_FPKM 10,965.1), which encodes the Juruin and Juruin_2. The arrows indicate the regions that correspond to each molecule. The cysteines that form disulfide bonds are highlighted in black.

**Appendice 6:**
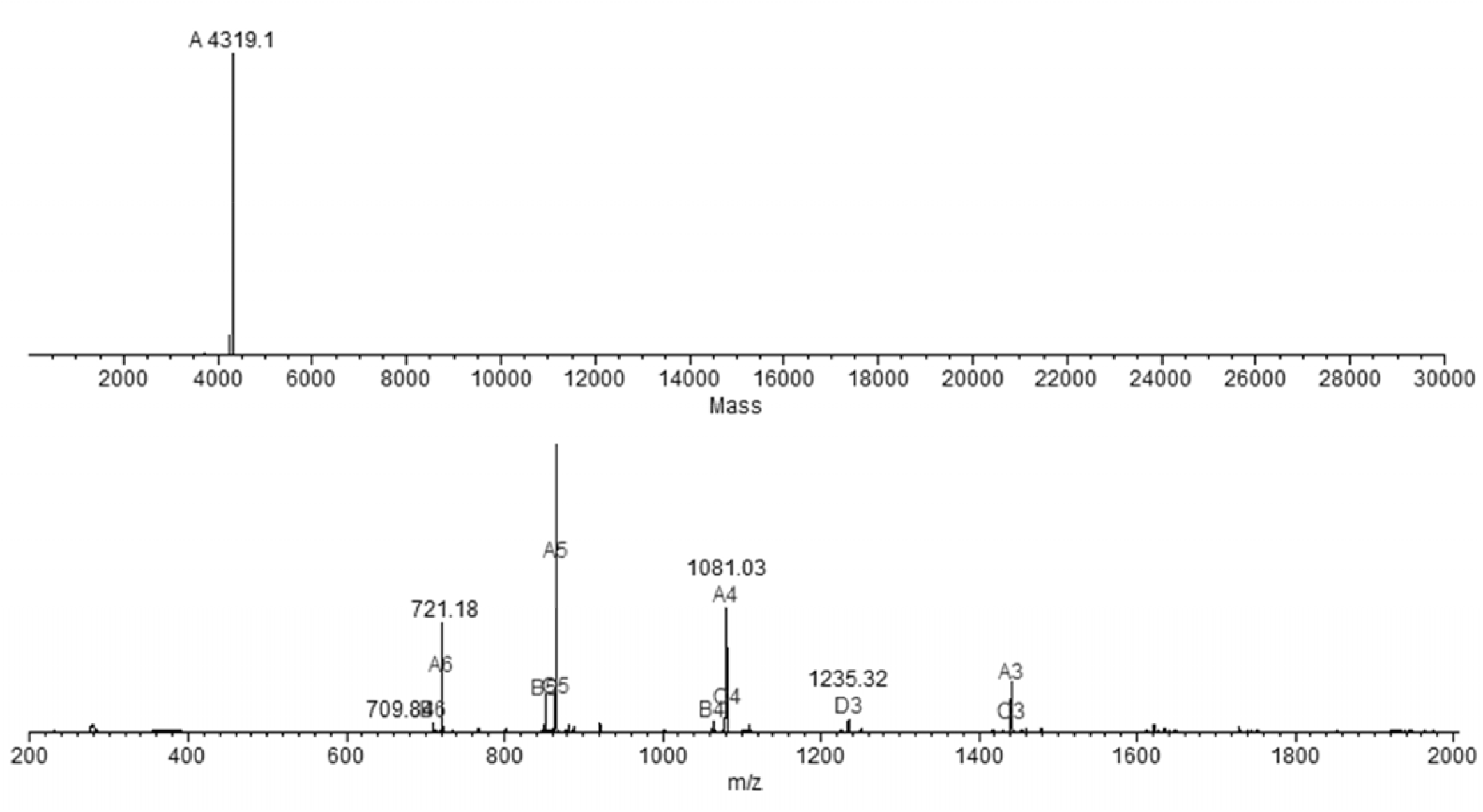
Determination of molecular mass of Juruin_2 by MassAnalyzer 1.03 (Amgen), the analysis indicated mass of 4,319.1 Da.

**Appendice 7:**
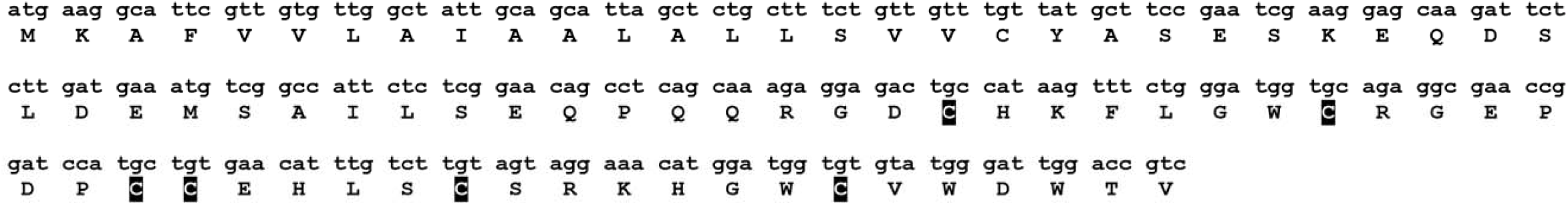
Nucleotide sequence and translated amino acids of NODE_17940 (Xpression_FPKM 460.63), which encodes the Juruenine. The cysteines that form disulfide bonds are highlighted in black.

**Appendice 8:**
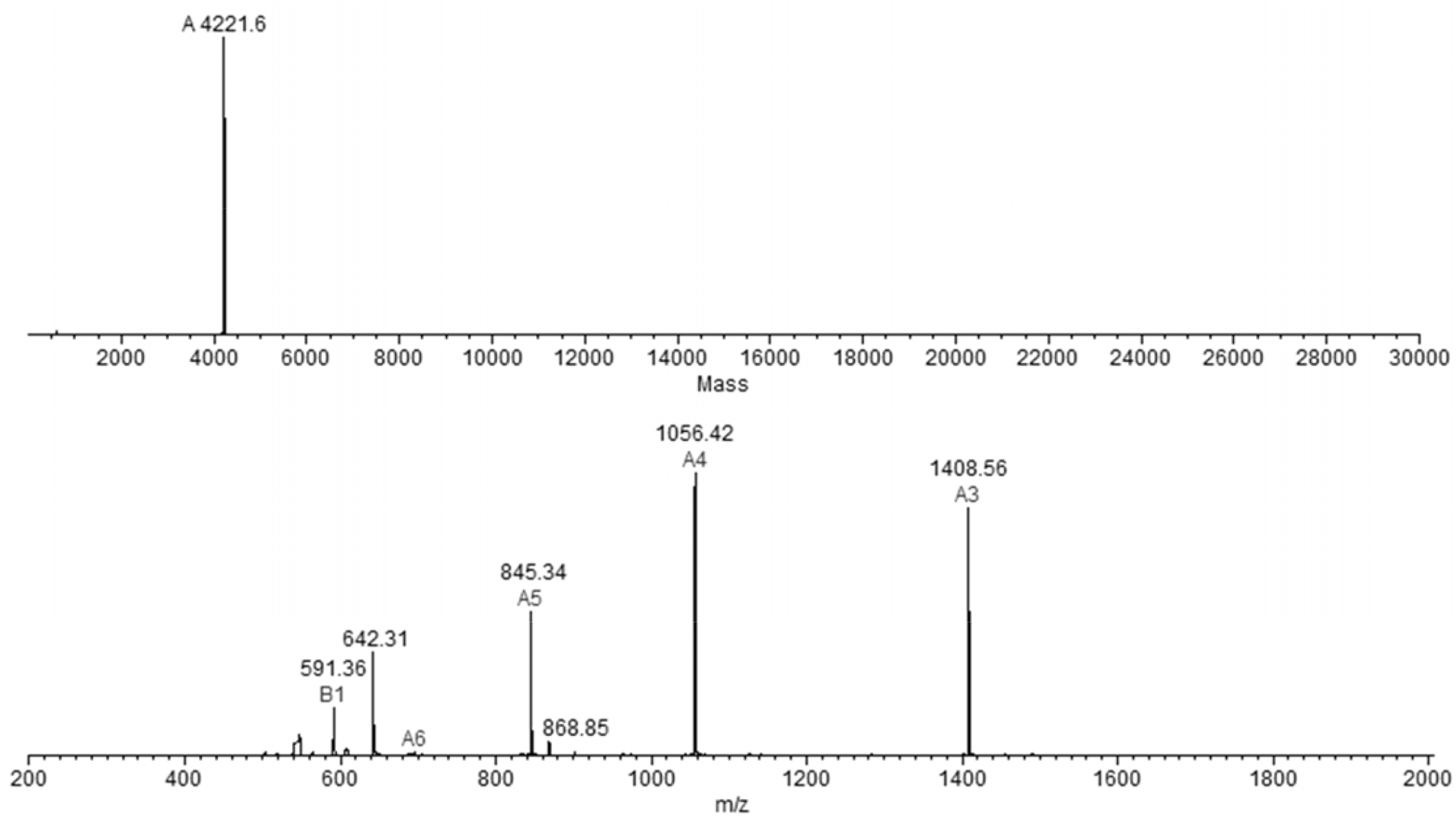
Determination of molecular mass of Juruenine by MassAnalyzer 1.03 (Amgen), the analysis indicated mass of 4,221.6 Da.

## Notes

### Competing Interest Statement

The authors have declared no competing interest.

